# Gut microbiome associations with poliovirus vaccine seroconversion in children in the Democratic Republic of the Congo

**DOI:** 10.1101/2025.10.13.682144

**Authors:** Jennifer A. Fulcher, Nicole A. Hoff, Fan Li, Patrick Mukadi, Kamy Musene, Sue K. Gerber, Megan Halbrook, Guillaume Ngoie Mwamba, Michel Kabamba, Nicole H. Tobin, Jean Jacques Muyembe-Tamfum, Grace M. Aldrovandi, Anne W. Rimoin

**Author notes:** Authors contributed equally (Jennifer A. Fulcher, Nicole A. Hoff, Grace M. Aldrovandi, Anne W. Rimoin).

## Abstract

**Background:** Despite efforts to increase global immunization, vaccination seroconversion in low and middle income countries (LMICs) is often lower than in high income countries (HICs). The reasons for this disparity are not fully understood. Given the role of the gut microbiome in immune development, we investigated the relationship between the gut microbiome and polio vaccine seroconversion in children in the Democratic Republic of the Congo (DRC).

**Methods:** This cross-sectional analysis included children ages 6-24 months old (n=90) enrolled in the DRC. Vaccine history was obtained from health records and/or self-report and poliovirus serostatus surveyed using dried blood spots. Nutritional status was evaluated using anthropomorphic measures. Microbiome profiling (16S rRNA gene) was performed and associations with poliovirus serostatus and malnutrition were examined.

**Results:** The average age of the study population was 13.6 months (SD=5.6) with 58% female. Poliovirus seropositivity was 65.5% and 22% of the children were malnourished. We found that presence of *Campylobacter* and *Veillonella*, especially at an early age, was associated with low poliovirus vaccination seroconversion. These bacterial taxa differed from those associated with malnutrition.

**Conclusions:** The presence of enteropathogens such as *Campylobacter* at a young age could be an important factor contributing to low vaccination seroconversion in children in LMICs.

## INTRODUCTION

Vaccine development has been one of the most significant scientific achievements in modern history. To date, effective vaccines exist to control or prevent 25 diseases[1]. When distributed effectively, vaccines have the potential to prevent millions of cases of permanent disability or death[2]. Despite these incredible achievements, there are an estimated 20 million under-vaccinated children across the globe with the majority residing in low and middle income countries (LMIC)[3]. In addition to challenges with distribution, oral vaccine responses in LMICs are less successful than in high income country (HIC) counterparts[4]. The reasons for the disparity in oral vaccination responses are not clear.

The commensal, symbiotic microbes that live in our gut, termed the microbiome, influence numerous physiological processes including metabolism, human disease, and immunity[5]. The development of a healthy gut microbiome in early childhood is critical for proper immune development[6]. In early infancy the gut microbiome composition is primarily influenced by the length of pregnancy, mode of delivery, and mode of feeding (breast milk, formula, or solid food)[7]. Other postnatal factors that alter the microbiome are antibiotics, environmental exposure, host genetics, and supplement use[8].

Not surprisingly, the microbiome also plays a role in the immune response to vaccinations, specifically oral vaccines[4, 9]. Infant responses to rotavirus vaccination in Pakistan correlate with increased Clostridium and Proteobacteria[10], and a similar study in Ghana showed rotavirus vaccine responses associate with increased *Streptococcus bovis* and decreased Bacteroidetes[11]. Studies using gnotobiotic pigs and fecal microbial transplant further showed that the gut microbiome alters rotavirus vaccine response[12], and that early administration of probiotics can improve vaccine response[13].

Associations with gut bacteria have also been observed with other oral vaccines. Increased *Bifidobacterium longum* correlated with T cell responses to live-attenuated oral polio vaccine (OPV), Bacillus Calmette-Guérin (BCG) and tetanus toxoid vaccines in infants in Bangladesh[14].

Near global eradication of poliovirus is one of the greatest vaccine triumphs. Both OPV and an injectable inactivated polio vaccine (IPV) are approved for use in preventing poliovirus infection.

Widespread distribution of OPV has effectively eradicated wild poliovirus types 2 and 3 while type 1 continues to circulate in only two countries worldwide[15]. However, despite the aggressive OPV campaigns in many LMICs, serological responses to poliovirus vaccines are suboptimal[16, 17]. This underimmunization contributes to the emergence of circulating vaccine-derived poliovirus (cVDPV), which are vaccine polio strains that have reacquired neurovirulence. cVDPV outbreaks have occurred in many countries, with the greatest number of cases in Nigeria and the Democratic Republic of the Congo (DRC)[18].

In DRC the routine polio immunization schedule for children includes four doses of bivalent OPV (at birth, 6, 10, and 14 weeks of age) and one dose of IPV (at 14 weeks of age). Additional supplemental monovalent vaccinations are distributed when cVDPV outbreaks occur. As most cVDPV are type 2, the monovalent OPV type 2 (mOPV2) has been used for these supplemental immunization activities (SIAs) in DRC[19]. A 2016 study analyzing population immunity to poliovirus in DRC found that among children aged 6-59 months 81% had immunity to type 1, 90% to type 2, and 70% to type 3[20]. However, a follow-up survey in 2018 found markedly decreased seroresponses with 43.8% to type 1, 41.1% to type 2, and 38% to type 3[21]. The mechanism for this diminished immunity is unknown; however, inadequate immunity has previously been correlated with age, wealth index, mother’s education, and urbanicity[20]. While certain gut bacteria have been associated with promoting OPV responses in Bangladesh[14], another study of children in India found little impact of gut bacteria on OPV responses but observed negative effects of non-polio enteric viruses on OPV response[22]. The relationship between the gut microbiome and OPV seroconversion remains unclear, but potentially important given the disparities in responses contributing to the emergence of cVDPV.

In this study we sought to examine relationships between the gut microbiome and polio vaccine serological responses in three close proximity geographic regions in DRC. We hypothesized that the gut microbiome composition would differ among these regions and influence polio vaccine seroconversion in children from DRC.

## METHODS

### Ethics Statement

This study was approved by the Institutional Review Board at the University of California, Los Angeles (UCLA IRB #18-000303) and the Ethics Committee at the Kinshasa School of Public Health in DRC. All participants provided informed consent prior to study entry. Parents or guardians provided consent for all children participants. Informed consent was administered in French, Swahili, or Kiluba by study administrators.

### Study Design and Participants

This study was a cross-sectional analysis using specimens and clinical data collected as part of a population-based serosurvey conducted in March 2018 among children aged 6-59 months old and their parent or guardian in DRC. Inclusion criteria for this analysis was children aged 6-24 months and willingness to provide oral and rectal swab samples. Approximately 10% of the larger ongoing serosurvey population was included in this analysis. Geographic regions for inclusion in this study were selected based on prior OPV SIAs and history of cVDPV outbreaks. The first region (Haut Lomami) included four health zones in Haut Lomami province which had nine cVDPV2 cases in the year prior to study collection and four or five (depending on health zone) mOPV2 SIAs prior to the study. The next region (Ankoro and Manono) was health zones in Tanganyika province which had ten cVDPV cases and two mOPV2 SIAs prior to the study. The third region (Kongolo and Kabalo) was health zones in Tanganyika province which had no cVDPV cases and no SIAs. Five villages were visited within each health zone. Villages were randomly selected using ArcGIS software (v10.5) ‘Create Random Point’ with the criteria: not in the same administrative Health Area and minimum separating distance of 500 meters. All houses in each selected village were sampled.

### Study Data and Specimens

Demographic and clinical data were collected from all participants after an informed consent process using tablets in the local language by trained interviewers. Dried blood spots (DBS) were collected via finger prick for serological assays. Rectal swabs and oral swabs (FLOQ swabs, Copan Diagnostics, Murrieta, CA) were collected by trained interviewers. All samples were collected on ice then stored at -20°C in Kinshasa, DRC before shipment to the USA. Once in the USA, rectal swabs were stored at -80°C until batch processing.

### Anthropomorphic Measures

Height, weight, and mid-upper arm circumference (MUAC) were measured from all participants using standard procedures. Z-scores based on the WHO Child Growth Standards were calculated using the ‘zscorer’ package in R(v4.1.2). To identify children with malnutrition we used both MUAC and weight-for-height (WFH) criteria to avoid excluding malnourished children that may be missed with either criterium alone[23]. Children with WFH Z-score less than or equal to -2 and/or MUAC less than or equal to 12.5cm were considered malnourished (wasting). We did not include stunting (height-for-age) or underweight (weight-for-age) in this analysis.

### Poliovirus Serologic Testing

DBS samples were shipped to the CDC Atlanta where neutralizing antibodies against poliovirus serotypes 1, 2, and 3 were quantified using a modified poliovirus microneutralization assay as previously described[20]. Neutralizing titers greater than or equal to 1:8 (3.0 log2) was used as the threshold for positive. In this study a positive serology for any poliovirus serotype was considered seropositive.

### DNA Extraction and 16S rRNA Gene Sequencing

Rectal swab specimens were transferred to Lysing Matrix E tubes (MP Biomedicals, Burlingame, CA) containing RLT lysis buffer (Qiagen, Hilden, Germany) and lysed using bead-beating on a TissueLyser (Qiagen). DNA was extracted using the AllPrep DNA/RNA/Protein kit (Qiagen) per the manufacturer’s protocol. Microbiome composition was analyzed by sequencing the V4 region of the 16S rRNA gene as previously described[24, 25]. Briefly, Golay-barcoded primers 515F/806R were used to amplify the V4 region in triplicate reactions. DNA extraction and PCR negative controls as well as bacterial mock community controls were included. PCR products were pooled and sequenced using an Illumina MiSeq with 2×150bp v2 chemistry.

### Microbiome and Statistical Analysis

Divisive Amplicon Denoising Algorithm (DADA2) version 1.19.1 was used for error correction, exact sequence inference, read merging, and chimera removal[26]. Taxonomic assignment was performed using RDP trainset 18[27] and contaminant sequences were removed using the ‘decontam’ R package (version 1.6.0). As previously described[28], we used inverse probability of treatment weighting to balance multiple confounders including age, sex, number of OPV doses, IPV status, geographic location, and other vaccinations. Permutational multivariate analysis of variance (PERMANOVA) as implemented in the ‘vegan’ R package (version 2.5-7) was used for assessing primary drivers of overall microbiome variation. Differential abundance testing was carried out using zero-inflated negative binomial models (‘pscl’ R package, version 1.5.5) with multiple testing correction by the Benjamini-Hochberg false discovery rate (FDR) method. Random forests classification models (‘ranger’ R package version 0.13.1) were utilized as an orthogonal approach to identify bacterial genera associated with vaccine seroconversion and malnutrition. Ten-fold cross-validation was used to identify the optimal number of genera in each model up to a maximum of 25 features to aid interpretability. One thousand forests each comprising 10,000 trees were used to obtain mean importance values. A sparse model was then constructed containing the selected number of features with the highest importance (calculated as mean decrease in accuracy). Model accuracy was calculated as percentage correctly classified from the out-of-bag error estimate. For both analyses, only genera present with at least 1% relative abundance in at least 10% of all samples were used for testing, resulting in 30 genera tested. All statistical analyses were performed using the R statistical environment v4.1.3. All sequence data is accessible via NCBI SRA under BioProject accession number PRJNA860001.

## RESULTS

### Study population

A total of 90 children from eight health zones in the Haut Lomami and Tanganyika provinces in DRC were included in the study (**Figure 1A)**. The study population ranged in age from 6 to 24 months (average 13.6 months, SD=5.6) with 42.2% males and 57.8% females.

**Figure 1.**
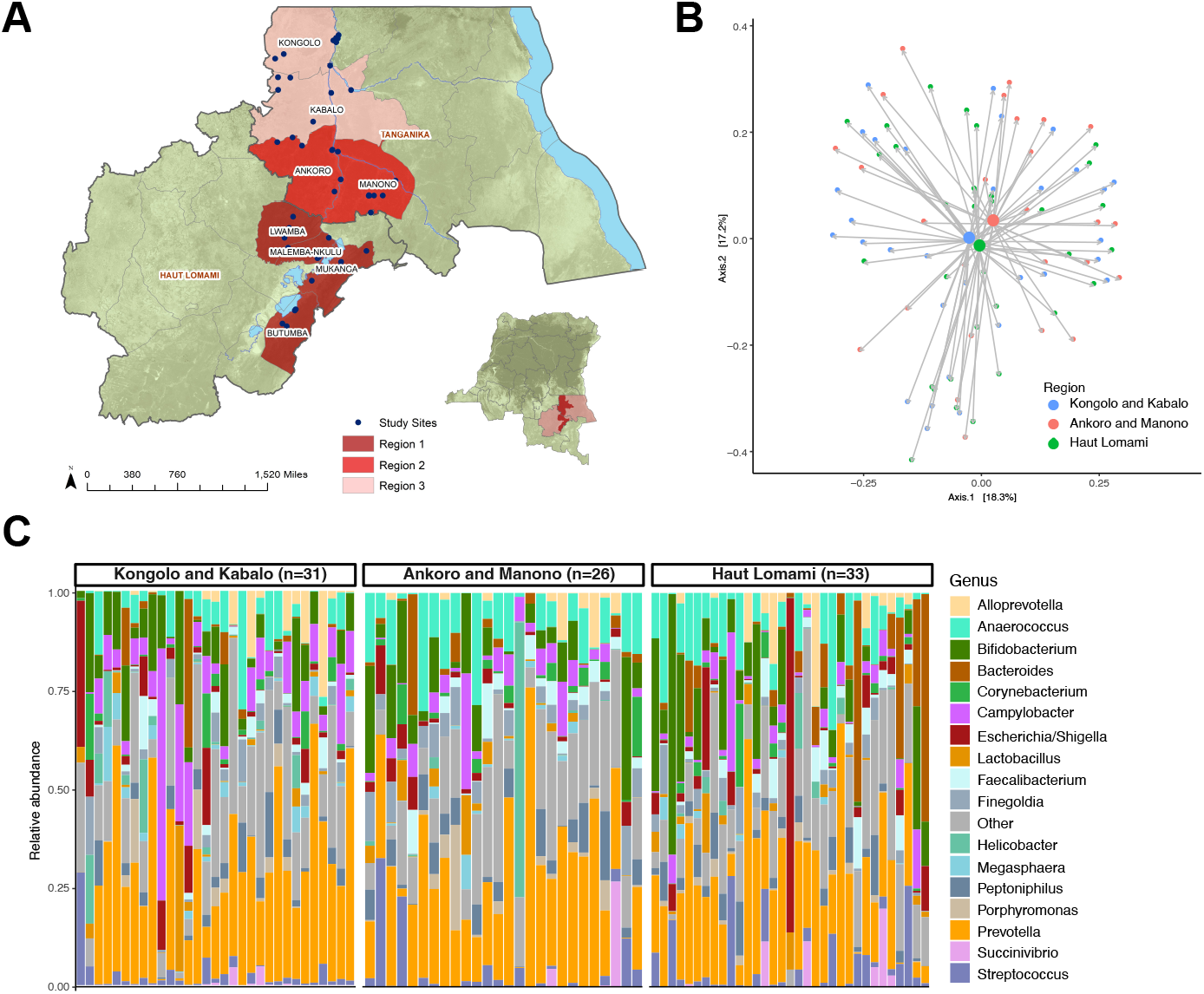
Gut microbiome in children across geographic regions in DRC. **(A)** Map of DRC showing the three geographic regions included in this study in the Haut Lomami and Tanganyika provinces. Each region included multiple health zones (labeled with names) and the villages visited in each region are denoted with blue dots. Regions 1 (Haut Lomami) and 2 (Ankoro and Manono) reported cVDPV cases and received supplemental immunization activities (SIAs) in the year preceding the study. Region 3 (Kongolo and Kabolo) did not report any cVDPV cases or receive any SIAs in the year preceding the study. **(B)** Principal coordinates analysis comparing geographic region using Jensen-Shannon divergence. **(C)** Taxa barplot showing microbiome composition grouped by geographic region. Individual bars within each group are ordered by participant age with youngest on left and oldest on right.

Demographic and vaccination history data is shown in **Table 1**. The diet is similar across all regions given the geographic proximity and similar terrain. The majority of infants in DRC are breastfed with similar rates across all regions.

**Table 1.** Study participant demographics and vaccination data.

|  | Geographic Region |  |  | p-value<sup>a</sup> |
| --- | --- | --- | --- | --- |
|  | Kongolo and Kabalo<br>n=31 | Ankoro and Manono<br>n=26 | Haut Lomami<br>n=33 |  |
| Age (months), mean (SD) | 12 (5.7) | 14 (5.9) | 15 (4.7) | 0.02 |
| Sex |  |  |  |  |
| Male, n (%) | 7 (22.6) | 13 (50) | 18 (54.5) | 0.02 |
| Female, n (%) | 24 (77.4) | 13 (50) | 15 (45.5) |  |
| Height (cm), mean (SD) | 68 (6.2) | 72 (7.2) | 72 (7.3) | 0.07 |
| Weight (kg), mean (SD) | 8.6 (1.8) | 9 (2.1) | 10 (2.9) | 0.10 |
| Mid-arm circumference (cm), mean (SD) | 14 (1.5) | 13 (1.4) | 14 (1.6) | 0.47 |
| Malnutrition, n (%) | 4 (13.8) | 10 (38.5) | 6 (18.8) | 0.07 |
| Immunization History<sup>b</sup> |  |  |  |  |
| Pentavalent<sup>c</sup>, n (%) | 19 (61.3) | 14 (58.3) | 15 (51.7) | 0.75 |
| Pneumococcus, n (%) | 18 (60) | 11 (47.8) | 13 (46.4) | 0.53 |
| BCG, n (%) | 17 (54.8) | 14 (56) | 17 (56.7) | 0.99 |
| Measles, n (%) | 12 (38.7) | 13 (52) | 12 (42.9) | 0.60 |
| Yellow Fever, n (%) | 12 (38.7) | 10 (40) | 11 (39.3) | >0.99 |
| Inactivated polio, n (%) | 13 (41.9) | 9 (37.5) | 11 (39.3) | 0.94 |
| Oral polio doses (routine)<sup>d</sup>, mean (SD) | 2.2 (1.8) | 2.2 (1.5) | 2 (1.9) | 0.90 |
| Oral polio doses (supplemental)<sup>e</sup>, mean (SD) | 0.55 (0.51) | 1.1 (1.1) | 2.4 (2.2) | <0.001 |
| Polio serology<sup>f</sup> |  |  |  |  |
| Type 1, n (%) | 16 (51.6) | 13 (50) | 17 (51.5) | >0.99 |
| Type 2, n (%) | 3 (9.7) | 14 (53.8) | 13 (39.4) | 0.001 |
| Type 3, n (%) | 14 (45.2) | 10 (38.5) | 11 (33.3) | 0.62 |
| Any type, n (%) | 19 (61.3) | 18 (69.2) | 22 (66.7) | 0.81 |

<sup>a</sup>p-values calculated using Chi-squared tests or Kruskal-Wallis tests<sup>b</sup>Immunization history obtained from vaccination cards as well as participant recall<sup>c</sup>Pentavalent vaccine includes diphtheria, pertussis, tetanus, Haemophilus influenzae, hepatitis B<sup>d</sup>Routine oral polio vaccine is bivalent type 1 and type 3<sup>e</sup>Supplemental oral polio vaccine is monovalent type 2<sup>f</sup>Performed at CDC; titers greater than 1:8 considered positive

### Geography and age are the biggest drivers of gut microbiome differences in children in DRC

PERMANOVA found age (p=0.007) and geographic region (p=0.044) to be the most significant factors in microbiome variation. The number of OPV doses also appeared to have greater influence (p=0.054) than other vaccination status (**Table 2**). The relative microbiome composition by geographic region is shown in **Figure 1B**, and comparison using principal coordinates analysis shown in **Figure 1C**.

**Table 2.** Factors contributing to gut microbiome variation in children in the DRC.

| Source of Variation | R<sup>2</sup> | p-value<sup>a</sup> |
| --- | --- | --- |
| Age | 0.016 | 0.007 |
| Sex | 0.009 | 0.133 |
| Geographic region | 0.021 | 0.044 |
| Oral polio vaccination (routine) | 0.004 | 0.888 |
| Oral polio vaccination (supplemental) | 0.012 | 0.054 |
| Inactivated polio vaccination | 0.005 | 0.746 |
| Bacille Calmette-Guerin (BCG) vaccination | 0.004 | 0.829 |
| Pentavalent vaccination<sup>b</sup> | 0.002 | 0.988 |
| Pneumococcus vaccination | 0.005 | 0.791 |
| Measles vaccination | 0.004 | 0.909 |
| Yellow Fever vaccination | 0.004 | 0.900 |
| Malnutrition | 0.006 | 0.540 |

<sup>a</sup>R<sup>2</sup> and p-values derived from permutational multivariate ANOVA using Jensen-Shannon divergence distances
<sup>b</sup>Pentavalent vaccine includes diphtheria, pertussis, tetanus, Haemophilus influenzae, hepatitis B

### Microbiome differences by polio vaccination seroconversion

We next analyzed the microbiome composition in children by polio vaccination response using poliovirus serology. The overall poliovirus seropositivity in our study population was similar across geographic regions with 61% in Kongolo and Kabalo health zones, 69% in Ankoro and Manono health zones, and 64% in selected health zones in Haut Lomami province. Since there was no major difference in seropositivity between geographic regions we used the full study cohort to examine the microbiome by poliovirus serostatus. There were no significant differences between children with positive or negative poliovirus serology in terms of age, sex, anthropomorphic measure, or other vaccination history (**Table 3**). We found increased alpha diversity **(Figure 2A**) and differences in overall composition when comparing children with polio vaccination seroconversion to those without (**Figures 2B and 2C**). To examine associations between specific bacteria and polio vaccination seroconversion we used zero-inflated negative binomial regression models with independent probability of treatment weighting adjustment for age, sex, geography, malnutrition, and other vaccinations. Decreased *Campylobacter* (p=0.002, FDR p_adj_=0.03) and *Veillonella* (p=0.0003, FDR p_adj_=0.005) were associated with a poliovirus vaccine seroconversion (**Figure 3A and Supplementary Table 1**). Random forests predictive modeling was also used to examine taxa associated with poliovirus vaccine seroconversion (**Figure 3B**). A sparse model comprising 19 features was selected by cross-validation and achieved an overall accuracy of 77.5% when classifying seropositive versus seronegative groups. Notably, *Campylobacter, Veillonella*, and also *Prevotella* were the most important taxa in distinguishing children with poliovirus vaccination seroconversion (**Figure 3C**). When looking at the relative abundance of these bacteria by child age, the seronegative group was characterized by presence of *Campylobacter* at early age (<12 months) and increased *Veillonella* across all ages (**Figure 3D**).

**Table 3.** Study participant demographics by poliovirus serostatus

|  | Poliovirus Serology<sup>a</sup> |  | p-value<sup>b</sup> |
| --- | --- | --- | --- |
|  | Negative<br>n=22 | Positive<br>n=58 |  |
| Age (months), mean (SD) | 13.2 (4.8) | 14.4 (5.8) | 0.46 |
| Sex |  |  | 0.41 |
| Male, n (%) | 8 (36%) | 27 (47%) |  |
| Female, n (%) | 14 (64%) | 31 (53%) |  |
| Height (cm), mean (SD) | 70.4 (8.2) | 71.7 (6.7) | 0.59 |
| Weight (kg), mean (SD) | 9.3 (2.6) | 9.5 (2.3) | 0.59 |
| Mid-arm circumference (cm), mean (SD) | 13.4 (2) | 14 (1.4) | 0.15 |
| Malnutrition, n (%) | 8 (36%) | 9 (16%) | 0.06 |
| Vaccination |  |  |  |
| Pentavalent, n (%) | 10 (45%) | 38 (66%) | 0.13 |
| Pneumococcus, n (%) | 8 (36%) | 34 (59%) | 0.09 |
| BCG, n (%) | 11 (50%) | 37 (64%) | 0.3 |
| Measles, n (%) | 8 (36%) | 29 (50%) | 0.32 |
| Yellow Fever, n (%) | 8 (36%) | 25 (43%) | 0.62 |
| Inactivated polio, n (%) | 6 (27%) | 27 (47%) | 0.14 |
| Oral polio doses (routine), mean (SD) | 1.8 (1.6) | 2.6 (1.6) | 0.08 |
| Oral polio doses (supplemental), mean (SD) | 1.8 (2.2) | 1.5 (1.5) | 0.98 |

<sup>a</sup>Positive serology includes positive for any poliovirus serotype; 10 participants were excluded due to no prior poliovirus vaccination
<sup>b</sup>p-values calculated using Fishers exact tests or Mann-Whitney tests

**Figure 2.**
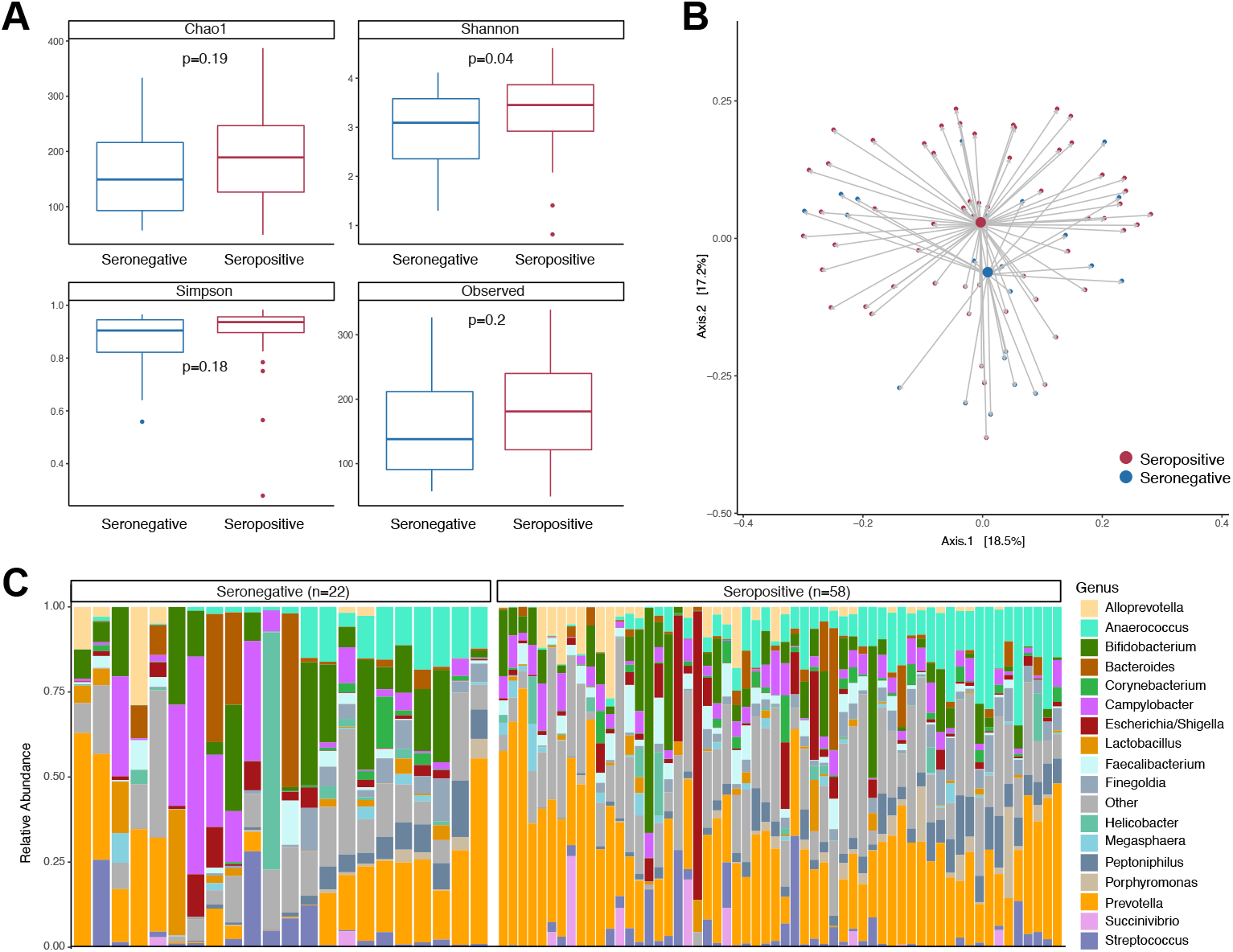
Microbiome differences by poliovirus serologic status. **(A)** Alpha diversity compared between poliovirus seropositive and seronegative groups. **(B)** Principal coordinates analysis comparing poliovirus seropositive and seronegative groups using Jensen-Shannon divergence. **(C)** Taxa barplot showing microbiome composition grouped by poliovirus serostatus. Individual bars within each group are ordered by participant age with youngest on left and oldest on right.

**Figure 3.**
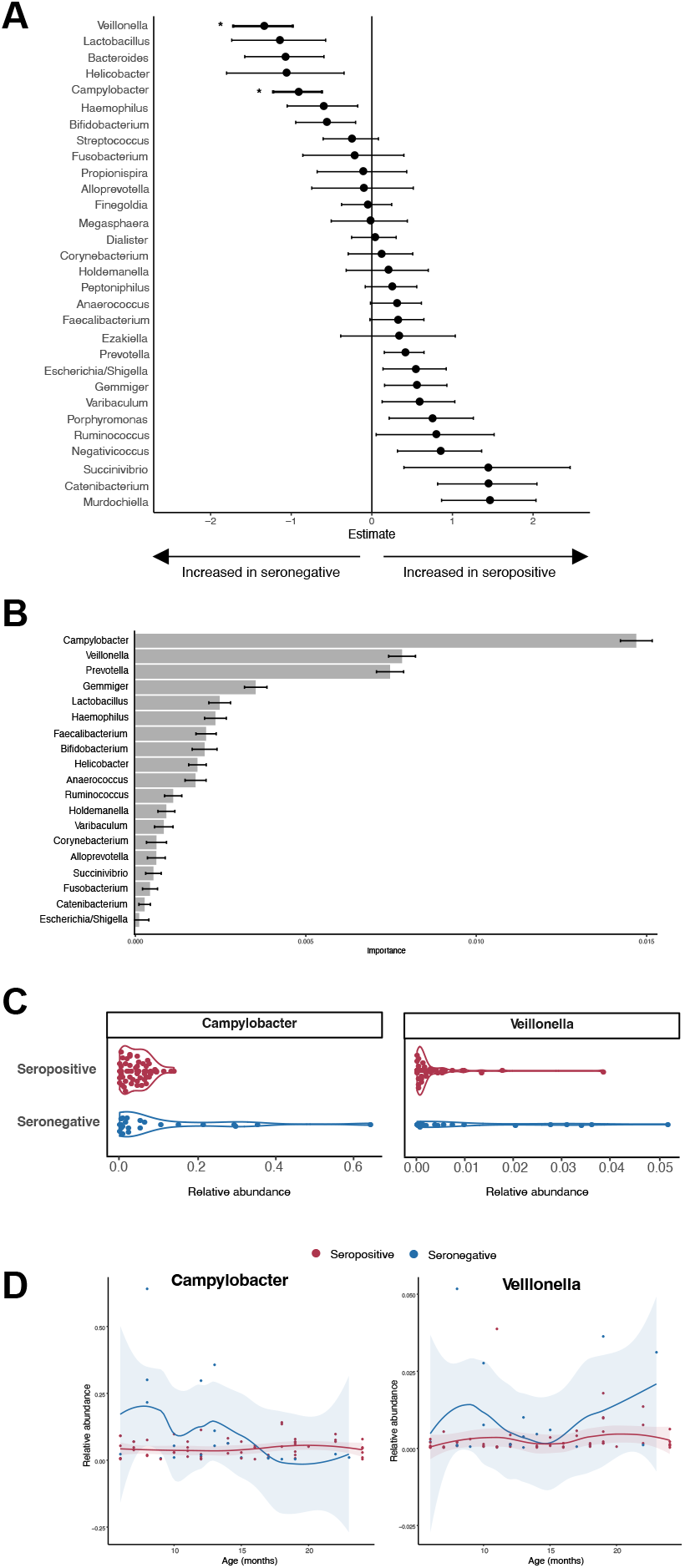
Bacterial taxa associated with poliovirus vaccination response. **(A)** Forest plot showing bacterial genera associated with positive poliovirus vaccination response. Values are shown as estimated coefficients and standard errors from a zero-inflated negative binomial regression model. Genera with an asterisk were statistically significant (FDR P_adj_<0.05). **(B)** Bacterial taxa that distinguish the microbiome of children with positive poliovirus vaccination response identified using random forests predictive modeling. Bacteria are ordered form most important feature (top) to least (bottom) with the relative importance shown on the x-axis. **(C and D)** Relative abundance of two most significant bacteria grouped by (C) seropositive (red) and seronegative (blue) and (D) ordered by age.

### Gut microbiome features in children with malnutrition differ from those characterizing low vaccine seroconversion

Since diet and nutrition are important contributors to the gut microbiome, we wanted to see if malnutrition could potentially explain the observed differences in gut microbiome and poliovirus vaccine seroconversion. Twenty-two percent of children in this study (n=20) met criteria for malnutrition using z-scores based on WHO growth standards. The poliovirus seropositivity was lower in the malnourished group (53%) compared to those without malnutrition (72%), suggesting that malnutrition could be a factor contributing to poor vaccine seroconversion. We compared the microbiome in children from all geographic regions with malnutrition compared to those without. Alpha diversity was decreased in the malnutrition group (**Figure 4A**), though the difference was not statistically significant. Differences in microbiome composition were observed in children with malnutrition (**Figure 4B and 4C**), with the most significant differences being less *Ruminococcus, Succinivibrio, Murdochiella*, and *Porphyromonas* in children with malnutrition (**Figure 4D and Supplementary Table 1**).

**Figure 4.**
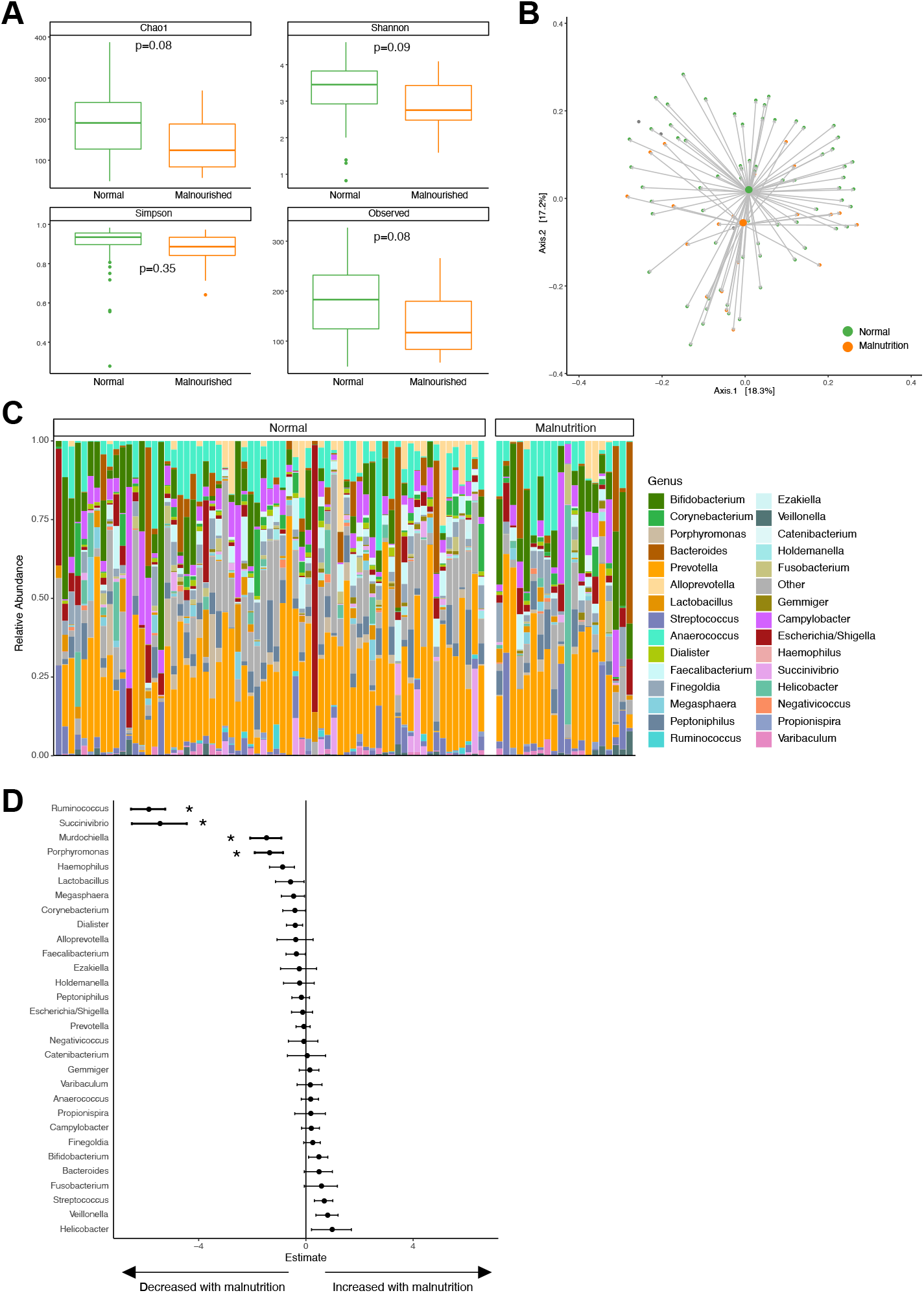
Microbiome differences by nutritional status. **(A)** Alpha diversity compared between children meeting malnutrition criteria (orange) and those who do not (green). **(B)** Principal coordinates analysis comparing children meeting malnutrition criteria (orange) and those who do not (green) using Jensen-Shannon divergence. **(C)** Taxa barplot showing microbiome composition grouped by nutritional status. Individual bars within each group are ordered by participant age with youngest on left and oldest on right. **(D)** Forest plot showing bacterial genera associated with malnutrition using zero-inflated negative binomial models. Values are shown as estimated coefficients and standard errors. Genera with an asterisk were statistically significant (FDR P_adj_<0.05).

## DISCUSSION

This study examined associations between the gut microbiome and poliovirus vaccination seroconversion in children in DRC. Geographic regions were selected based on cVDPV cases and SIAs, yet we did not find significant differences in poliovirus seropositivity and gut microbiome taxa between these regions (Figure 1). We did find that the presence of *Campylobacter* and *Veillonella*, especially at an early age, was associated with low poliovirus vaccine seroconversion when examining the total study population (Figure 3). Approximately 20% of the study population met criteria for malnutrition, and the bacterial taxa altered in the malnutrition group were distinct from those associated with poliovirus vaccination seroconversion (Figure 4).

The infant gut microbiome differs between HICs and LMICs, and these differences have been hypothesized to contribute to the disparate vaccination responses in these countries[4]. *Prevotella-*rich microbiomes are prevalent in many LMICs while microbiomes in HICs have less *Prevotella* and greater *Bacteroides*. Increases in this Prevotella/Bacteroides ratio has been negatively associated with rotavirus vaccination responses in Ghana[11]. The gut microbiome in our study population from DRC exhibited the characteristic *Prevotella* dominance seen in LMICs, consistent with prior studies in this region[29], however we did not find significant associations between Prevotella/Bacteroides and poliovirus vaccination seroconversion. Our findings also differ from at least two prior studies in that we did not find significant associations with *Bifidobacteria* and OPV response, though differences in study population age may at least partially explain these differences[14, 30]. The precise microbiome features that contribute to vaccination response likely vary by geography, host age/genetics, and vaccine antigen. Generalization across existing studies is further challenged by differences in study populations, sequencing technologies, analysis strategies, and study design. Thus, large coordinated multi-site studies are needed to ultimately answer these important questions.

The presence of *Campylobacter* was associated with low poliovirus vaccine seroconversion in our study. This is consistent with prior studies that found that the presence of enteropathogens, both bacterial and viral, has negative effects on vaccine response[31-33]. *Campylobacter* can be a common cause of diarrhea, and it is not known if this finding represents asymptomatic carriage or is a consequence of recent diarrheal infection in these children. Ongoing exposure to enteropathogens can lead to a subclinical gastrointestinal inflammation termed environmental enteropathy (EE). This chronic mucosal inflammation inhibits proper microbiome colonization and immune development, especially when occurring at early age[34]. Conflicting evidence exists regarding the relationship between EE and vaccination response[35], but several studies have found negative associations between EE and OPV seroconversion[31, 36, 37]. Perhaps the presence of *Campylobacter* in our study population contributes to inflammation and risk of EE leading to poor poliovirus vaccine seroconversion. The large MAL-ED birth cohort study conducted in eight LMICs similarly associated early *Campylobacter* exposure with poor OPV response[33], reinforcing the importance of reducing these exposures in early life. The presence of *Veillonella* in the gut can also signify inflammation. While *Veillonella* is typically a healthy colonizer, especially in the oral cavity, when found in the gut it can be pro-inflammatory thus adding to the unfavorable mucosal immune environment in these children[38].

Malnutrition has been implicated in poor vaccine responses in LMICs[4]. Approximately one-fifth of our study population met criteria for malnutrition. Other studies in DRC have reported higher malnutrition rates[39], though differences in time, location, and nutrition criteria (malnourished vs stunting) could account for these differences. Nutrition plays an important role in shaping the gut microbiome, thus malnutrition can have deleterious effects on microbiome composition[40]. Subramanian et al showed that malnutrition delays gut microbiome maturation in children in Bangladesh, with *Faecalibacterium prausnitzii* and *Ruminococcus* the most important taxa in distinguishing children with malnutrition[41]. We were unable to perform a similar analysis in our cohort due to the cross-sectional study design, but did identify *Ruminococcus* as significantly decreased in children with malnutrition. Of note, the taxa associated with poliovirus vaccine seroconversion and malnutrition were different, suggesting that influences beyond just malnutrition likely explain our findings.

There are several caveats to consider when interpreting these study results. First, the study size was modest and it is possible additional microbiome differences may have been identified with a larger sample size. Second, vaccination history data was obtained by either health records/vaccination cards (when available) or self-report, and therefore could be subject to recall and/or reporter bias. As vaccination history was used as a covariate this could influence our results. Finally, this study only assessed gut bacteria but did not examine enteric viruses or fungi. Enteric viruses have had reported associations with OPV response[22, 31, 42] and animal models indicate intestinal fungi influence immune responses[43]. There were no cases of cVDPV in our study population thus we could not directly investigate microbiome factors that may contribute to cVDPVs. As the geographic regions samples had different rates of cVDPVs we hypothesized we might indirectly identify potential microbiome factors relating to cVDPV prevalence; however, we did not find significant differences. Continued investigation into the relationship between the gut microbiome and cVDPV is warranted, especially given the known associations between the gut microbiome and poliovirus replication and pathogenesis[44].

This study adds to the growing literature supporting the potential role of the gut microbiome in vaccination seroconversion, especially in LMICs. Despite this, there remains unresolved questions regarding the most important bacteria affecting vaccine responses. Further research to better define these microbial factors, as well as how to promote a healthy microbiome, will be important to improve vaccination success globally.

## ACKNOWLEDGEMENTS

We are grateful to all participants in this study for their generous participation. We would like to acknowledge the contributions of the late Dr. Emile Okitolonda-Wemakoy to this work. We also thank Dr. William Weldon and the Poliovirus Laboratory at the Centers for Disease Control and Prevention (CDC) in Atlanta, GA for assistance with poliovirus serology and Katharine Newman (UCLA) for research assistance.

## REFERENCES

1. World Health Organization. Global vaccine action plan 2011-2020. Available at: https://www.who.int/publications/i/item/global-vaccine-action-plan-2011-2020. Accessed July 1 2022.

2. Ehreth J. The value of vaccination: a global perspective. Vaccine 2003; 21:4105–17.

3. UNICEF. UNICEF Data: Immunization. Available at: https://data.unicef.org/topic/child-health/immunization/. Accessed July 1 2022.

4. Lynn DJ, Benson SC, Lynn MA, Pulendran B. Modulation of immune responses to vaccination by the microbiota: implications and potential mechanisms. Nat Rev Immunol 2021.

5. Lynch SV, Pedersen O. The Human Intestinal Microbiome in Health and Disease. N Engl J Med 2016; 375:2369-79.

6. Belkaid Y, Hand TW. Role of the microbiota in immunity and inflammation. Cell 2014; 157:121–41.

7. Backhed F, Roswall J, Peng Y, et al. Dynamics and Stabilization of the Human Gut Microbiome during the First Year of Life. Cell Host Microbe 2015; 17:690–703.

8. Tamburini S, Shen N, Wu HC, Clemente JC. The microbiome in early life: implications for health outcomes. Nat Med 2016; 22:713–22.

9. Zimmermann P, Curtis N. The influence of the intestinal microbiome on vaccine responses. Vaccine 2018; 36:4433–9.

10. Harris V, Ali A, Fuentes S, et al. Rotavirus vaccine response correlates with the infant gut microbiota composition in Pakistan. Gut Microbes 2017:1–9.

11. Harris VC, Armah G, Fuentes S, et al. Significant Correlation Between the Infant Gut Microbiome and Rotavirus Vaccine Response in Rural Ghana. J Infect Dis 2017; 215:34–41.

12. Twitchell EL, Tin C, Wen K, et al. Modeling human enteric dysbiosis and rotavirus immunity in gnotobiotic pigs. Gut Pathogens 2016; 8:51.

13. Chattha KS, Vlasova AN, Kandasamy S, Rajashekara G, Saif LJ. Divergent immunomodulating effects of probiotics on T cell responses to oral attenuated human rotavirus vaccine and virulent human rotavirus infection in a neonatal gnotobiotic piglet disease model. J Immunol 2013; 191:2446–56.

14. Huda MN, Lewis Z, Kalanetra KM, et al. Stool microbiota and vaccine responses of infants. Pediatrics 2014; 134:e362–72.

15. Global Polio Eradication Initiative. Endemic Countries. Available at: https://polioeradication.org/where-we-work/polio-endemic-countries/. Accessed June 5 2022.

16. Patriarca PA, Wright PF, John TJ. Factors affecting the immunogenicity of oral poliovirus vaccine in developing countries: review. Rev Infect Dis 1991; 13:926–39.

17. Okayasu H, Sutter RW, Czerkinsky C, Ogra PL. Mucosal immunity and poliovirus vaccines: impact on wild poliovirus infection and transmission. Vaccine 2011; 29:8205-14.

18. Global Polio Eradication Initiative. Circulating Vaccine Derived Poliovirus. Available at: http://polioeradication.org/polio-today/polio-now/this-week/circulating-vaccine-derived-poliovirus/. Accessed June 5 2022.

19. Mbaeyi C, Alleman MM, Ehrhardt D, et al. Update on Vaccine-Derived Poliovirus Outbreaks - Democratic Republic of the Congo and Horn of Africa, 2017-2018. MMWR Morb Mortal Wkly Rep 2019; 68:225–30.

20. Voorman A, Hoff NA, Doshi RH, et al. Polio immunity and the impact of mass immunization campaigns in the Democratic Republic of the Congo. Vaccine 2017; 35:5693–9.

21. Halbrook M, Gadoth A, Mukadi P, et al. Poliovirus-Neutralizing Antibody Seroprevalence and Vaccine Habits in a Vaccine-Derived Poliovirus Outbreak Region in the Democratic Republic of Congo in 2018: The Impact on the Global Eradication Initiative. Vaccines (Basel) 2024; 12.

22. Praharaj I, Parker EPK, Giri S, et al. Influence of Nonpolio Enteroviruses and the Bacterial Gut Microbiota on Oral Poliovirus Vaccine Response: A Study from South India. J Infect Dis 2019; 219:1178–86.

23. Schwinger C, Golden MH, Grellety E, Roberfroid D, Guesdon B. Severe acute malnutrition and mortality in children in the community: Comparison of indicators in a multi-country pooled analysis. PLoS One 2019; 14:e0219745.

24. Bender JM, Li F, Martelly S, et al. Maternal HIV infection influences the microbiome of HIV-uninfected infants. Sci Transl Med 2016; 8:349ra100.

25. Pannaraj PS, Li F, Cerini C, et al. Association Between Breast Milk Bacterial Communities and Establishment and Development of the Infant Gut Microbiome. JAMA Pediatr 2017; 171:647–54.

26. Callahan BJ, McMurdie PJ, Rosen MJ, Han AW, Johnson AJ, Holmes SP. DADA2: High-resolution sample inference from Illumina amplicon data. Nat Methods 2016; 13:581–3.

27. Wang Q, Garrity GM, Tiedje JM, Cole JR. Naive Bayesian classifier for rapid assignment of rRNA sequences into the new bacterial taxonomy. Appl Environ Microbiol 2007; 73:5261–7.

28. Cook RR, Fulcher JA, Tobin NH, et al. Effects of HIV viremia on the gastrointestinal microbiome of young MSM. Aids 2019; 33:793–804.

29. Brazier L, Elguero E, Koumavor CK, et al. Evolution in fecal bacterial/viral composition in infants of two central African countries (Gabon and Republic of the Congo) during their first month of life. PLoS One 2017; 12:e0185569.

30. Zhao T, Li J, Fu Y, et al. Influence of gut microbiota on mucosal IgA antibody response to the polio vaccine. NPJ Vaccines 2020; 5:47.

31. Grassly NC, Praharaj I, Babji S, et al. The effect of azithromycin on the immunogenicity of oral poliovirus vaccine: a double-blind randomised placebo-controlled trial in seronegative Indian infants. Lancet Infect Dis 2016; 16:905–14.

32. Myaux JA, Unicomb L, Besser RE, et al. Effect of diarrhea on the humoral response to oral polio vaccination. Pediatr Infect Dis J 1996; 15:204–9.

33. Pan WK, Seidman JC, Ali A, et al. Oral polio vaccine response in the MAL-ED birth cohort study: Considerations for polio eradication strategies. Vaccine 2019; 37:352–65.

34. Ordiz MI, Stephenson K, Agapova S, et al. Environmental Enteric Dysfunction and the Fecal Microbiota in Malawian Children. Am J Trop Med Hyg 2017; 96:473–6.

35. Church JA, Parker EP, Kosek MN, et al. Exploring the relationship between environmental enteric dysfunction and oral vaccine responses. Future Microbiol 2018; 13:1055–70.

36. Kosek MN, Mduma E, Kosek PS, et al. Plasma Tryptophan and the Kynurenine-Tryptophan Ratio are Associated with the Acquisition of Statural Growth Deficits and Oral Vaccine Underperformance in Populations with Environmental Enteropathy. Am J Trop Med Hyg 2016; 95:928–37.

37. Naylor C, Lu M, Haque R, et al. Environmental Enteropathy, Oral Vaccine Failure and Growth Faltering in Infants in Bangladesh. EBioMedicine 2015; 2:1759–66.

38. Read E, Curtis MA, Neves JF. The role of oral bacteria in inflammatory bowel disease. Nat Rev Gastroenterol Hepatol 2021; 18:731–42.

39. Kandala NB, Madungu TP, Emina JB, Nzita KP, Cappuccio FP. Malnutrition among children under the age of five in the Democratic Republic of Congo (DRC): does geographic location matter? BMC Public Health 2011; 11:261.

40. Kane AV, Dinh DM, Ward HD. Childhood malnutrition and the intestinal microbiome. Pediatr Res 2015; 77:256–62.

41. Subramanian S, Huq S, Yatsunenko T, et al. Persistent gut microbiota immaturity in malnourished Bangladeshi children. Nature 2014; 510:417–21.

42. Parker EP, Kampmann B, Kang G, Grassly NC. Influence of enteric infections on response to oral poliovirus vaccine: a systematic review and meta-analysis. J Infect Dis 2014; 210:853–64.

43. Wheeler ML, Limon JJ, Bar AS, et al. Immunological Consequences of Intestinal Fungal Dysbiosis. Cell Host Microbe 2016; 19:865–73.

44. Kuss SK, Best GT, Etheredge CA, et al. Intestinal microbiota promote enteric virus replication and systemic pathogenesis. Science 2011; 334:249–52.

